# Genetic background shapes SEZ6L2 autoimmunity and reveals coordinated immune responses linked to neurological dysfunction

**DOI:** 10.64898/2026.03.31.715689

**Authors:** Carlos J. Reyes-Sepúlveda, John Randolph, Julia M. Granato, Alison Hobbins, Jennetta W. Hammond

## Abstract

SEZ6L2 autoantibodies have been identified in patients with subacute cerebellar ataxia, but the underlying immune mechanisms and pathogenic pathways remain poorly understood. We previously established a C57BL/6 mouse model of SEZ6L2 autoimmunity that recapitulates key features of the disease. Here, we evaluated whether genetic background influences the magnitude and organization of SEZ6L2-directed immune responses. Pilot screening of autoimmune-prone strains identified SJL mice as exhibiting accelerated and enhanced antibody responses following SEZ6L2 immunization. In a large-cohort study, SEZ6L2-immunized SJL mice developed robust and sustained antibody responses, along with antigen-specific CD4⁺ and CD8⁺ T-cell activation. Expanded immune profiling revealed increased CNS infiltration of multiple lymphocyte populations, including CD4⁺ T cells, CD8⁺ T cells, B cells, and dendritic cells, as well as the presence of SEZ6L2-specific B cells within the brain. In addition, SJL mice exhibited strain-specific immunodominant T-cell epitopes distinct from those observed in C57BL/6 mice. Functionally, SEZ6L2-immunized SJL mice developed motor deficits consistent with cerebellar dysfunction. Integration of behavioral outcomes demonstrated a consistent overall impairment, and multivariate analysis revealed that coordinated humoral and cellular immune responses were associated with behavioral deficits. Together, these findings demonstrate that SEZ6L2-directed immune responses produce coordinated adaptive immune activation linked to neurological dysfunction and establish the SJL strain as an enhanced model for studying SEZ6L2 autoimmunity. This model also provides a platform for investigating disease mechanisms and therapeutic strategies.

## INTRODUCTION

Immune-mediated cerebellar disorders represent a significant but often underrecognized cause of progressive neurological disability.^1, 2^ Many patients present with unexplained ataxia in which immune involvement becomes evident only after detection of disease-associated neural autoantibodies. The presence of specific neural autoantibodies can provide insight into disease mechanisms, prognosis, and therapeutic responsiveness. However, for many recently identified neuronal autoantigens, including SEZ6L2, the immune pathways driving disease and their relationship to neurological dysfunction remain unclear.

Autoantibodies targeting seizure-related protein 6-like 2 (SEZ6L2) have been identified in patients with subacute cerebellar ataxia characterized by progressive gait, instability, dysarthria, ocular motor abnormalities, and cognitive symptoms.^3–16^ Neuroimaging frequently demonstrates cerebellar atrophy with limited overt inflammatory findings. So far, responses to immunotherapy have been variable, with some patients exhibiting stabilization or partial improvement of symptoms.^3–14^ Since their initial description, reported cases remain limited and diagnostic testing is still not widely available. Although SEZ6L2 autoantibodies appear to be disease-specific, ^3, 5^ it is not yet known whether they are directly pathogenic or how they interact with cellular immune responses to drive neurological dysfunction.

SEZ6L2 is a transmembrane protein broadly expressed in neurons and neuroendocrine cells, with the cerebellum exhibiting the highest protein levels.^17, 18^ Human genetic studies have linked *Sez6l2* variants to autism spectrum disorder, Alzheimer’s disease, and various autoimmune conditions.^19–30^ Although SEZ6L2’s molecular functions in neural circuits remain incompletely defined, proposed roles include regulation of complement activity and trafficking proteins such as GLUA1, ADD2, and cathepsin D. ^5, 31–35^ *Sez6l2* knockout mice exhibit progressive motor impairment, gait abnormalities, reduced grip strength, and cognitive deficits.^18, 36, 37^ At the synaptic level, loss of SEZ6L2 is associated with reduced dendritic spine length and decreased synaptic protein expression, indicative of impaired synaptic connectivity.^36^ These phenotypes parallel those observed in SEZ6L2-immunized mice,^17^ and support a loss-of-function disease mechanism.

In the present study, we sought to determine whether genetic background influences the magnitude and organization of SEZ6L2-directed immune responses and whether this could provide an enhanced platform for mechanistic investigation. Through pilot screening of autoimmune-prone strains, we identified SJL mice as exhibiting accelerated and amplified immune responses following SEZ6L2 immunization. We therefore performed a large-cohort study to characterize humoral and cellular immunity, define antigen-specific T- and B-cell responses, and assess neurological outcomes.

Our findings demonstrate that SEZ6L2 immunization in SJL mice induces coordinated humoral and cellular immune activation, expanded CNS immune infiltration, and measurable motor deficits. In addition, this model enables identification of antigen-specific lymphocyte populations and reveals a quantitative relationship between immune responses and neurological dysfunction. Together, these results establish the SJL strain as an enhanced and mechanistically informative model for studying SEZ6L2-directed autoimmunity.

## MATERIALS AND METHODS

### General Animal

Animal care and use were carried out in compliance with the US National Research Council’s Guide for the Care and Use of Laboratory Animals and the US Public Health Service’s Policy on Humane Care and Use of Laboratory Animals. Protocols were approved by the University Committee on Animal Resources at the University of Rochester.

All mice were obtained from The Jackson Laboratory and maintained under standard housing conditions with 3-5 mice per cage and ad libitum access to food and water. Animals were randomly assigned to experimental groups, and investigators were blinded to immunization status during behavioral and immunological analyses. Male and female mixed sex cohorts were used for C57BL/6, SWR, Balb/c, and DBA/1 strains. For the SJL strain, only female mice were used, as male mice are extremely aggressive and cannot be socially housed. Additionally, our previous study with C57BL/6 suggested that there were no sex differences in response to SEZ6L2 immunization.^17^

### SEZ6L2 Immunization Protocol and experimental timelines

Immunization was performed using our established protocol with the extracellular domain (1-742) of human SEZ6L2 (GenBank: BC000567; Protein ID: AAH00567.1; encoding the splice variant that is 809 amino acids in length).^17^ Briefly, mice were immunized subcutaneously with recombinant human SEZ6L2 emulsified in Complete Freund’s Adjuvant with final emulsion concentrations of 5mg/mL SEZ6L2 and 2mg/mL Mycobacterium tuberculosis. Two dosing regimens were used: low dose (50 μL for delivery of 250 μg SEZ6L2 and 100 μg heat-inactivated Mycobacterium tuberculosis) and high dose (200 μL for delivery of 1 mg SEZ6L2 and 400 μg Mycobacterium tuberculosis). Each animal received 1 or 4 dorsal site injections of 50 μL each. Sham control mice were injected with 200 μL of adjuvant alone, without SEZ6L2. To induce CNS autoimmunity, pertussis toxin (250 ng; Hooke Laboratory, BT-0105) was administered intraperitoneally on days 0 and 1. Serum was collected at three weeks post-immunization, followed by a booster immunization with SEZ6L2 (1mg/mL; 100μL) in Incomplete Freund’s Adjuvant and repeated pertussis toxin administration. At six weeks post-immunization, mice were euthanized, and serum, spleen, and brain tissue were collected for downstream analyses.

#### Strain test cohort

For each strain, five mice (12-13 weeks of age) were immunized with the high dose of SEZ6L2 and compared with two sham-immunized controls as well as the other immunized strains. Because experiments were conducted across two independent days, two cohorts of C57BL/6 mice were included (n = 4–5 per cohort; 9 total) to provide an internal reference.

#### Large SJL cohort

The large SJL/J cohort was immunized at 8 weeks of age with either the low or high dose of SEZ6L2 (named SEZ6L2^Lo^ and SEZ6L2^Hi^ respectively).

### Behavioral Assessments

#### Open field, wire grid foot fault, and accelerating rotarod

assays were conducted as previously described.^17^ The Rout (Q=1%) method was used to check for outliers in these datasets; however, none were found or removed.

#### Tapered beam walk

Mice were evaluated for motor coordination using a tapered beam walk test. The beam measured 2″ in width at the start, narrowing to 3/16″ at the distal end, with a secondary platform approximately 1/2″ below the walking surface to provide side ledges in case of slips. Mice were habituated to the apparatus for at least 10 minutes the day prior to testing and completed two practice runs immediately before two recorded trials. Foot faults were quantified during traversal, while time to cross was also measured; however, no group differences were observed for the latter metric.

#### Pole test

Mice were placed head-down at the top of a 50–60 cm vertical metal pole (∼1 cm diameter) with textured grooves to facilitate grip. The time required to descend to the base without falling was recorded. Outliers were identified using the ROUT method (Q = 1%), resulting in the exclusion of three mice (1 sham, 2 SEZ6L2^Lo^) that had prolonged pauses in their descent.

#### Novel Object Recognition

During habituation, mice were placed in an empty arena for 10 minutes; this session also served as the open field assessment. The following day, mice were exposed to two identical objects (smooth doorknobs) for 5 minutes. After a 4-hour interval, mice were reintroduced to the arena containing one familiar object and one novel object matched for size but differing in shape and texture. Exploration was recorded for 5 minutes. Time spent interacting with each object was quantified, and a novel object preference index was calculated as: (time with novel − time with familiar) / total object interaction time. Outliers were identified using the ROUT method (Q = 1%), and three mice were excluded (1 SHAM, 2 SEZ6L2^Hi^).

#### Integrative Behavioral z-score

To facilitate interpretation across behavioral assays, we applied Z-normalization and calculated an integrated behavioral Z-score. For each assay, Z-scores were computed as the SEZ6L2-immunized mean minus the sham mean divided by the sham standard deviation. To ensure consistent directionality across measures, scores were oriented such that higher Z-values reflect worse performance. For assays in which impairment corresponded to negative values, Z-scores were inverted (multiplied by −1) to align directionality across all metrics. The standardized scores were then summed to generate a composite measure of overall behavioral impairment.

### Serum antibody ELISA

Anti-SEZ6L2 IgG levels were quantified using an ELISA assay as previously reported.^17^ Briefly, plates were coated with recombinant mouse SEZ6L2 protein (1ng/µl in 100µl PBS), incubated with diluted serum samples (1:1000), and developed using horseradish peroxidase–conjugated anti-mouse IgGs (Bio-Rad, 1706516, 1:15,000), TMB substrate (ThermoScientific, N301), and 0.18 M sulfuric acid stop solution. Absorbance was measured using a SpectraMax M5 plate reader.

### Splenocyte isolation and *in vitro* stimulation with SEZ6L2 recombinant protein or peptides for IFNγ ELISPOT and flow cytometry assays

Spleens were collected and processed for ELISPOT and flow cytometry assays as previously described.^17^ For the ELISPOT assays, 350,000 splenocytes were plated onto IFNγ ELISPOT plates from either Cellular Technology Limited (CTL; mIFNgp-2M) or BD Biosciences (mouse IFNγ ELISPOT kit) and stimulated with vehicle, recombinant mouse SEZ6L2 protein (8μg/mL), or individual peptides (10μg/mL) in serum-free CTL media for 24 hours. ELISPOT plates were then processed according to the manufacturer’s instructions and spots were counted using an ELISPOT reader (ImmunoSpot 5.2.12; Cellular Technology Limited). Peptide stimulation was performed for T-cell epitope mapping using the same SEZ6L2 15-mer peptide library with a 10 amino acid overlap that was previously used with C57BL6/J mice.^17^ The library includes peptides spanning the extracellular domains of both human and mouse SEZ6L2 proteins.

For surface and intracellular cytokine staining followed by flow cytometry analysis, splenocytes were plated at two million cells per well in a U-bottom 96-well plate with serum-free CTL media supplemented with 20 μg/mL mouse recombinant SEZ6L2 protein. The cells were incubated at 37°C for either 18 or 68 hours. Then cells were treated with Brefeldin A (BioLegend, 420601; 1:1000) and incubated for another 6 hours. The cells were then processed and stained for surface lineage markers and intracellular cytokines using the flow panel in Table 1, prior to flow cytometric acquisition with a BD Symphony A1 cytometer.

**Table 1:**
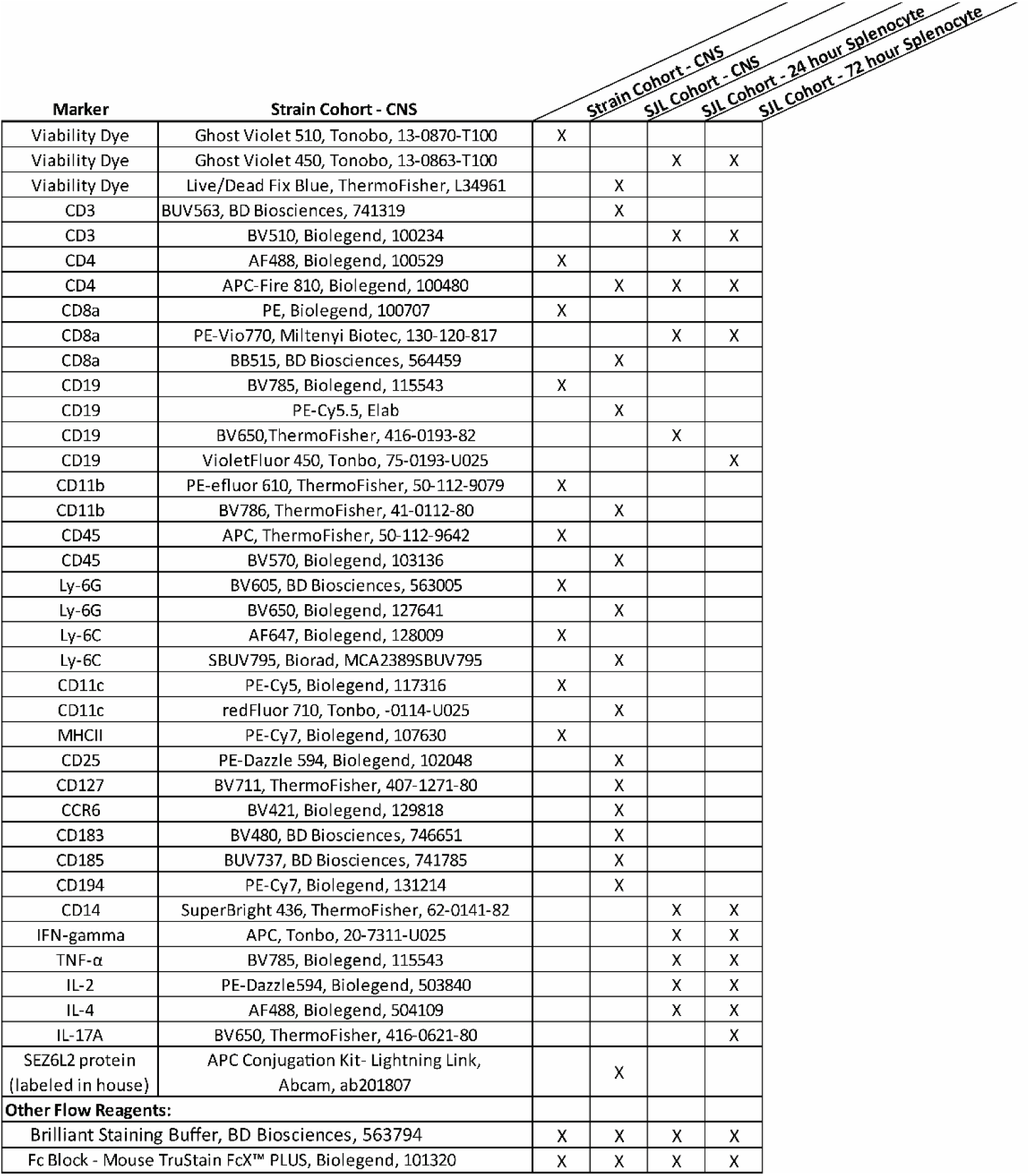
Antibodies and other reagents used for flow cytometry assays.

### Isolation and flow cytometry analysis of brain-infiltrating leukocytes

Brain mononuclear cells were isolated following transcardial perfusion. For the strain test cohort, minced brains were incubated with collagenase D (Roche, 11088866001) and DNase (Roche, 11088866001) for 30 minutes at 37°C followed by Percoll density gradient separation as described previously.^17^ For the large SJL cohort, brains were instead homogenized in cold RPMI media with 10mM HEPES, 5% FBS, and 100μg/mL DNase without additional enzymatic digestion. After filtering through a 70μm filter, isotonic Percoll was added to the cell suspensions to a final concentration of 38% isotonic Percoll. The mixtures were spun at 500xg for 30 minutes at 4°C. For both cohorts, the cell pellets were washed 2x in FACS buffer (PBS with 1% BSA, 25μg/mL DNase, and 1mM EGTA) prior to cell surface staining as previously described with the flow panels in Table 1. To detect SEZ6L2-specific B cells in the CNS, 75μg of the human SEZ6L2 extracellular domain was fluorescently labeled according to kit instructions with the APC Conjugation Kit- Lightning Link (Abcam, cat # ab201807). SEZ6L2-APC was incubated with cells along with the flow panel antibodies. The strain cohort was analyzed using a BD LSR II Flow cytometer, whereas the SJL cohort was analyzed with a Cytek Aurora full-spectrum cytometer. Data was analyzed by FlowJo^TM^ Software (v10.10.0_CL).

### Statistical Analysis

Statistical analyses were performed using GraphPad Prism software (versions 10.3.1 and11.0.0, La Jolla California, USA). Data were assessed for normality and lognormality using the Shapiro–Wilk and D’Agostino–Pearson tests. Variance was evaluated using Brown–Forsythe and Bartlett’s tests. Additional statistical tests and sample sizes for individual datasets are specified in the figure legends, with *n* representing individual animals. Data are presented as the mean ± SEM on a linear scale or as the geometric mean ± SD on a log scale. A threshold of p < 0.05 was considered statistically significant and is indicated on graphs by the statistical markers: *p<0.05, **p<0.01, ***p<0.001.

## Results

### Pilot strain screening identifies SJL mice as enhanced humoral responders to SEZ6L2 immunization with increased CNS leukocyte infiltration

To assess whether genetic background influences SEZ6L2-directed autoimmune responses, we immunized five autoimmune-prone mouse strains (C57BL/6, SWR, BALB/c, DBA/1, and SJL)^38, 39^ with recombinant human SEZ6L2 (Figure 1A). All strains generated detectable antibody responses against mouse SEZ6L2 following immunization. Notably, SJL mice exhibited significantly elevated anti-SEZ6L2 IgG levels by three weeks post-immunization compared to all other strains. By six weeks post-immunization, antibody levels in most strains increased to levels comparable to those observed in SJL mice at the earlier time point, whereas DBA/1 mice showed a comparatively weaker response (Figure 1B). These results indicate that the SJL background supports accelerated and enhanced humoral responses capable of recognizing endogenous SEZ6L2.

**Figure 1:**
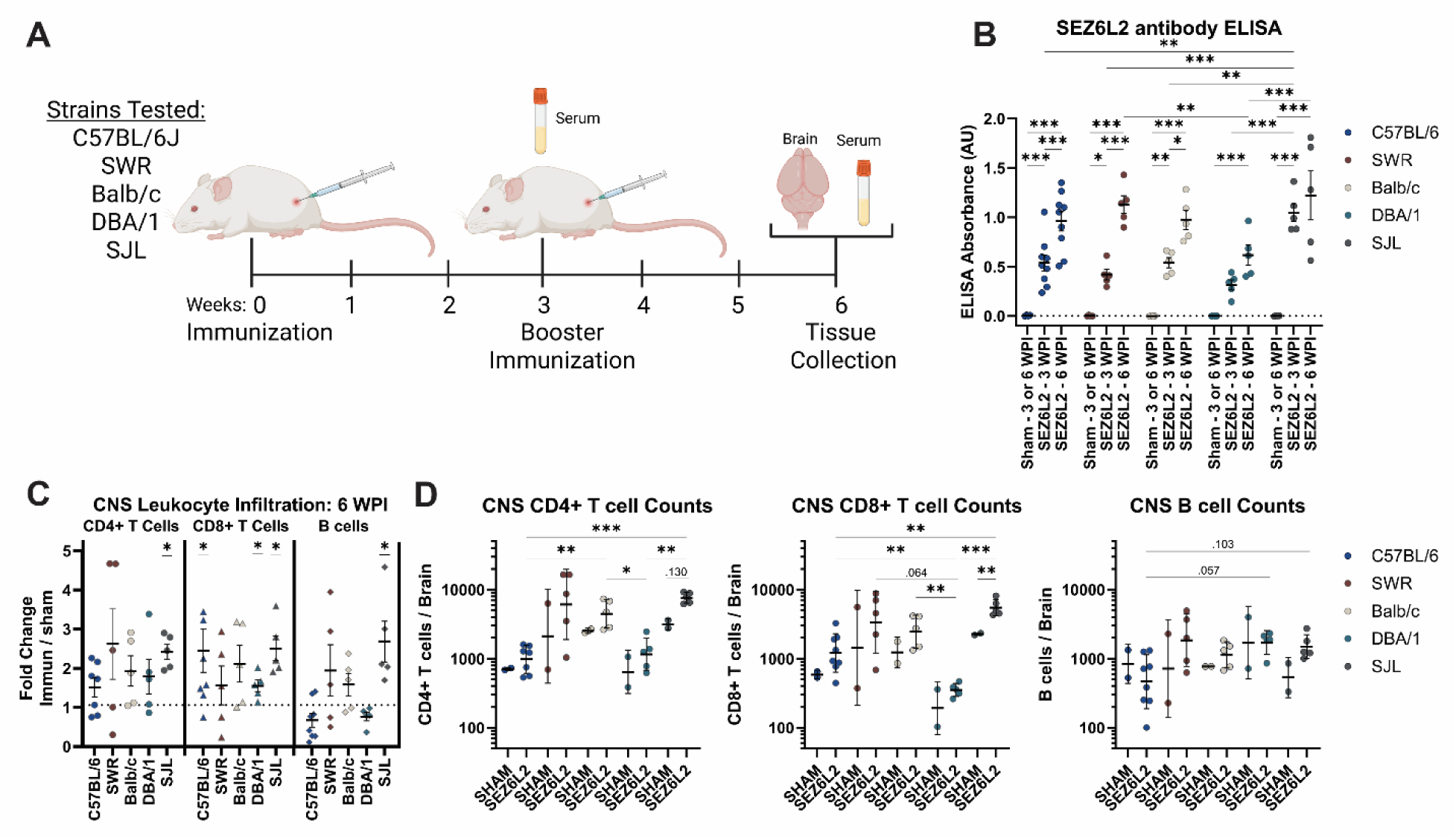
SEZ6L2 immunization induces strain-dependent humoral responses and CNS immune cell infiltration. **A)** Experimental timeline. **B)** Serum anti-SEZ6L2 IgG levels measured by ELISA at 3- and 6-weeks post-immunization (WPI). Statistics: 2-way ANOVA with Tukey’s multiple comparisons test; Data are presented as mean +/- SE. **C)** Fold change in CNS leukocyte infiltration in SEZ6L2-immunized mice relative to strain-matched sham controls. Statistics: one sample t-test compared to a theoretical mean of 1; Data are presented as mean +/- SE. **D)** Absolute counts of CD4⁺ T cells, CD8⁺ T cells, and CD19⁺ B cells in the CNS. Statistics: Welch’s ANOVA with Dunnett’s T3 multiple comparisons test; Data are presented as geometric mean +/- SD. **For all graphs:** Each point represents an individual mouse. Sham: n = 2 mice per strain; SEZ6L2-immunized: n = 5 mice per strain (except for C57BL/6, n = 9).

We next evaluated CNS immune infiltration by quantifying CD4⁺ T cells, CD8⁺ T cells, and CD19⁺ B cells (Figure 1C–D). Across strains, SEZ6L2 immunization produced trends toward increased T-cell infiltration relative to sham controls. In contrast, SJL mice demonstrated robust and statistically significant increases in CD4⁺ T cells, CD8⁺ T cells, and B cells, with each leukocyte population elevated by approximately 2.5-fold compared to strain-matched controls. Consistent with this, SJL mice exhibited the highest overall CNS T-cell counts among all strains tested. Together, these findings indicate that the SJL background supports an amplified cellular immune response within the CNS following SEZ6L2 immunization.

Based on the combined enhancement of humoral responses and CNS immune infiltration, we selected SJL mice for subsequent studies to determine whether this amplified immune profile is associated with neurological dysfunction and an ataxia-like phenotype.

### SEZ6L2 immunization induces robust and sustained antibody levels in SJL mice

To further characterize immune responses in SJL mice, we performed a large-cohort immunization study using either a low or high dose of recombinant human SEZ6L2, along with sham-immunized controls (n = 21 mice per group; Figure 2A). Serum ELISA analysis demonstrated robust anti-SEZ6L2 IgG production in both low- and high-dose groups at 3-and 6-weeks post-immunization (Figure 2B). Consistent with the pilot cohort (Figure 1), antibody levels were already elevated at 3 weeks and remained sustained at 6 weeks. Although the high-dose group exhibited a trend towards increased antibody levels compared to the low-dose group at both timepoints, this difference did not reach statistical significance (p = 0.19 at both timepoints). Because ELISA plates were coated with recombinant mouse SEZ6L2, these data confirm that the antibody response is cross-reactive with endogenous antigen. Together, these findings reinforce that SJL mice mount a rapid and sustained humoral response following SEZ6L2 immunization.

**Figure 2:**
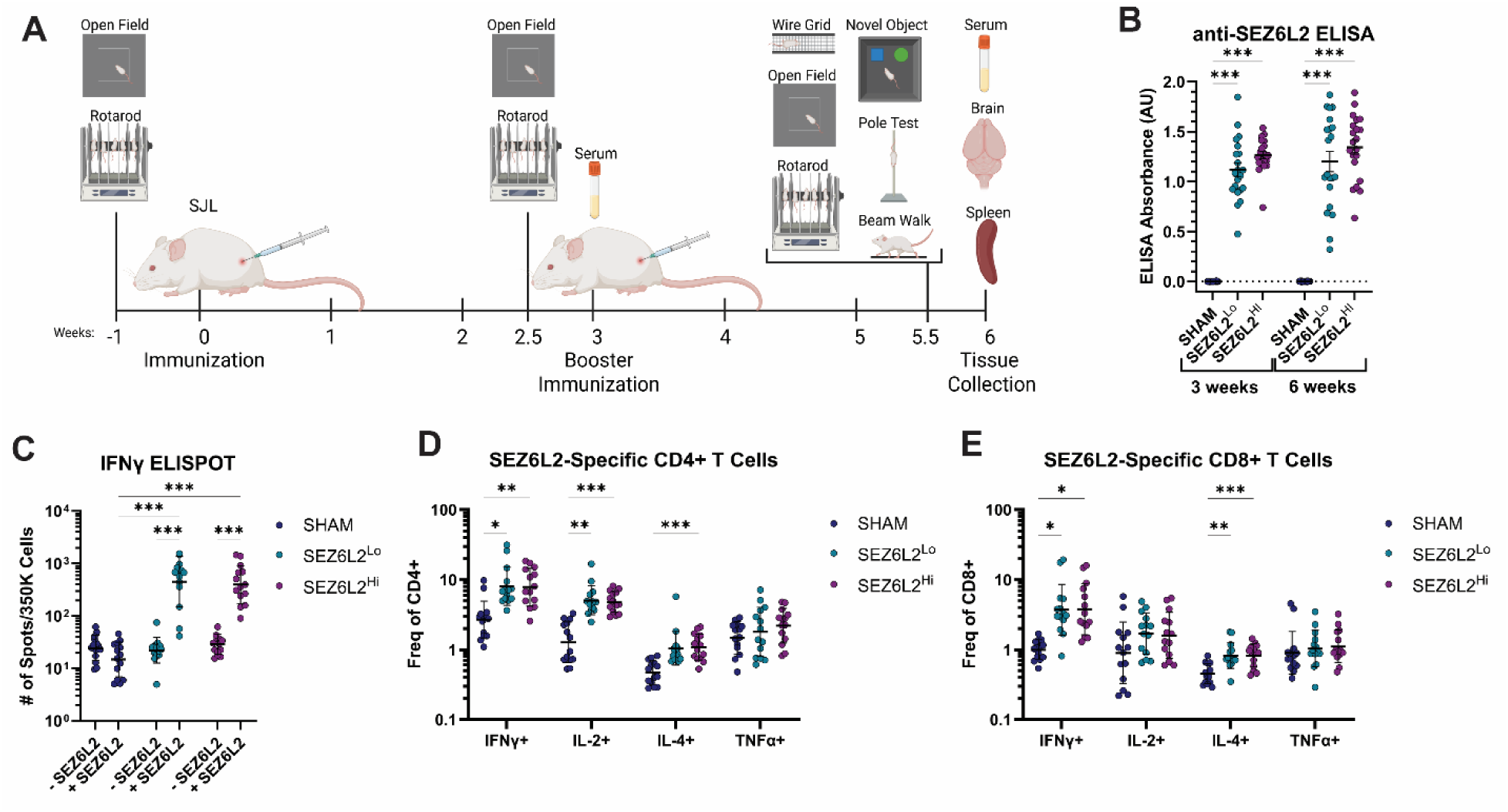
SEZ6L2 immunization induces robust humoral and antigen-specific T-cell responses in SJL mice. **A)** Experimental timeline for the large SJL cohort. Female mice were immunized at 8 weeks of age with either a low (Lo) or high (Hi) dose of recombinant SEZ6L2. **B)** Serum anti-SEZ6L2 IgG levels measured by ELISA at 3- and 6- weeks post-immunization (WPI). Statistics: repeated-measures 2-Way ANOVA with Tukey’s multiple comparisons test. Data are presented as mean +/- SE; n=21-22 mice per group. **C)** IFNγ ELISPOT analysis of splenocytes from sham- and SEZ6L2-immunized mice following stimulation with recombinant mouse SEZ6L2. Statistics: repeated-measures 2-Way ANOVA with Tukey’s multiple comparisons test; Data are presented as geometric means +/- SD; n=14 mice per group. **D-E)** Flow cytometric analysis of the intracellular cytokine production in CD4⁺ (D) and CD8⁺ (E) T cells following stimulation with recombinant SEZ6L2. Statistics: repeated-measures 2-Way ANOVA with Geisser-Greenhouse correction and Tukey’s multiple comparisons test; Data are presented as geometric means +/- SD; n=14 mice per group.

### SEZ6L2 immunization elicits antigen-specific CD4⁺ and CD8⁺ T-cell responses

To assess antigen-specific T-cell responses, splenocytes from immunized and control mice were stimulated with recombinant mouse SEZ6L2 and analyzed by IFNγ ELISPOT. SEZ6L2-immunized mice exhibited a significant increase in IFNγ-producing cells in response to antigen stimulation compared to sham controls (Figure 2C). No significant differences were observed between low- and high-dose groups, indicating that both immunization regimens generate comparable antigen-specific T-cell responses.

We next performed intracellular cytokine staining and flow cytometric analysis to further characterize T-cell polarization. SEZ6L2 immunization resulted in significant increases in the frequency of IFNγ⁺ and IL-17^+^ CD4⁺ and CD8⁺ T cells, consistent with a TH1/TH17-skewed response (Figure 2D–E; Supplemental Figure 2). This was further supported by an increase in IL-2–producing CD4⁺ T cells. In contrast, TNFα-producing T-cell populations were not significantly altered. A modest but significant increase in IL-4 expressing CD4⁺ and CD8⁺ T cells was also observed, indicating the presence of a secondary TH2 component.

Together, these data demonstrate that SEZ6L2 immunization in SJL mice induces a robust pool of antigen-specific CD4⁺ and CD8⁺ T cells with a predominantly TH1/TH17 phenotype and a minor TH2 contribution, capable of recognizing endogenous SEZ6L2.

### SJL mice exhibit distinct immunodominant T-cell epitopes following SEZ6L2 immunization

To define SEZ6L2-specific T-cell epitopes, we utilized a 15-mer peptide library with 10 amino acid overlap spanning the extracellular domains of both human and mouse SEZ6L2 (188 total peptides). Because immunization was performed using human SEZ6L2, but the autoimmune response must be directed toward endogenous mouse SEZ6L2, this combined library enabled simultaneous assessment of both cross-reactive (“autoimmune”) and human-specific (“foreign”) epitopes within a single assay.

At 6 weeks post-immunization, splenocytes from SEZ6L2-immunized SJL mice were stimulated with individual peptides and analyzed by IFNγ ELISPOT (Figure 3). All mice exhibited strong reactivity to multiple peptides unique to the human SEZ6L2 sequence, primarily localized to the region spanning amino acids 41–90. This region corresponds to the least conserved portion of the SEZ6L2 extracellular domain between species (Supplementary Figure 3), which likely facilitates T-cell recognition of human-specific epitopes without the need to overcome central or peripheral tolerance mechanisms. Within this same region, we also identified a peptide unique to the mouse SEZ6L2 sequence that elicited a measurable response in approximately half of the immunized mice, suggesting partial crossover of the response toward endogenous epitopes. Additionally, two immunodominant peptides shared between human and mouse SEZ6L2 were identified at amino acids 491–505 (detected in 10/10 mice) and 581–595 (detected in 5/10 mice), demonstrating that conserved regions of the protein can also serve as endogenous epitopes. Finally, a few other peptides elicited low-frequency responses in one or two mice, particularly within the region spanning amino acids 611–705, indicating a broader but less consistent epitope repertoire.

**Figure 3:**
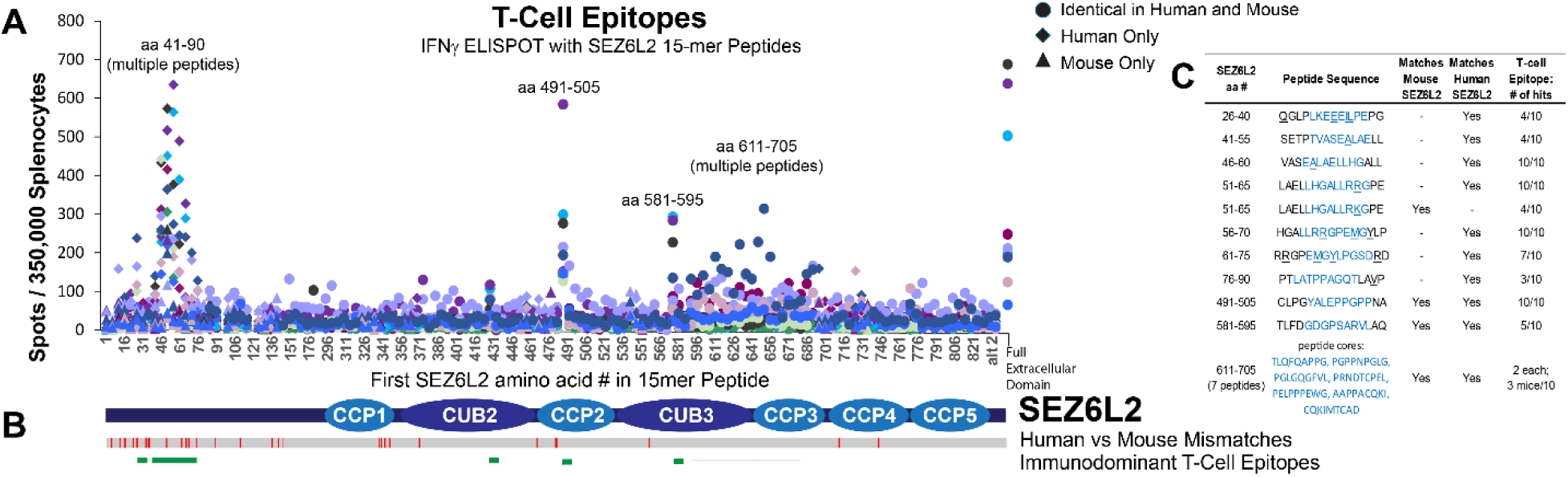
Identification of immunodominant SEZ6L2 T-cell epitopes in SJL mice. **A)** IFNγ ELISPOT analysis of splenocytes from SEZ6L2-immunized SJL mice stimulated with a 15-mer peptide library (10 amino acid overlap) spanning the extracellular domains of human and mouse SEZ6L2. Each point represents a single peptide response from an individual mouse (n = 10). Circles indicate peptides conserved between human and mouse, diamonds indicate human-specific peptides, and triangles indicate mouse-specific peptides. Colors denote individual mice. **B)** Schematic of SEZ6L2 domain structure showing regions of sequence divergence (red), immunodominant epitopes (green), and alignment with ELISPOT responses in panel A. **C)** Summary table of peptides eliciting positive ELISPOT responses. Predicted MHC class II (I-A^s^) core binding regions (IEDB; NetMHCIIpan 4.1 EL) are highlighted in blue. Underlined residues indicate sequence differences between human and mouse SEZ6L2 protein.

It is important to note that the immunodominant epitopes identified in SJL mice did not overlap with those previously observed in C57BL/6 mice.^17^ This shift in epitope hierarchy is consistent with differences in MHC class II haplotypes between strains (SJL: I-A^s^; C57BL/6: I-A^b^) and highlights the influence of host genetics on antigen presentation and T-cell specificity.

### SJL Mice Exhibit Expanded CNS Immune Infiltration Following SEZ6L2 Immunization

To determine whether enhanced humoral responses in SJL mice were associated with increased CNS immune infiltration, we performed flow cytometry on CD45⁺ brain leukocytes at 6 weeks post-immunization. Compared to sham controls, SEZ6L2-immunized SJL mice exhibited significant increases in CD4⁺ T cells, CD8⁺ T cells, CD19⁺ B cells, and CD11c⁺ dendritic cells (Figure 4A–B). This infiltration profile was broader than that observed in C57BL/6 mice, in which immune expansion was largely restricted to CD4⁺ T cells with minimal changes in CD8⁺ T-cell and B-cell populations.^17^ In contrast, most myeloid populations (including microglia, monocytes, and neutrophils) were unchanged or only modestly altered, indicating that immune expansion in SJL mice primarily reflects adaptive immune recruitment rather than global innate activation and infiltration.

**Figure 4:**
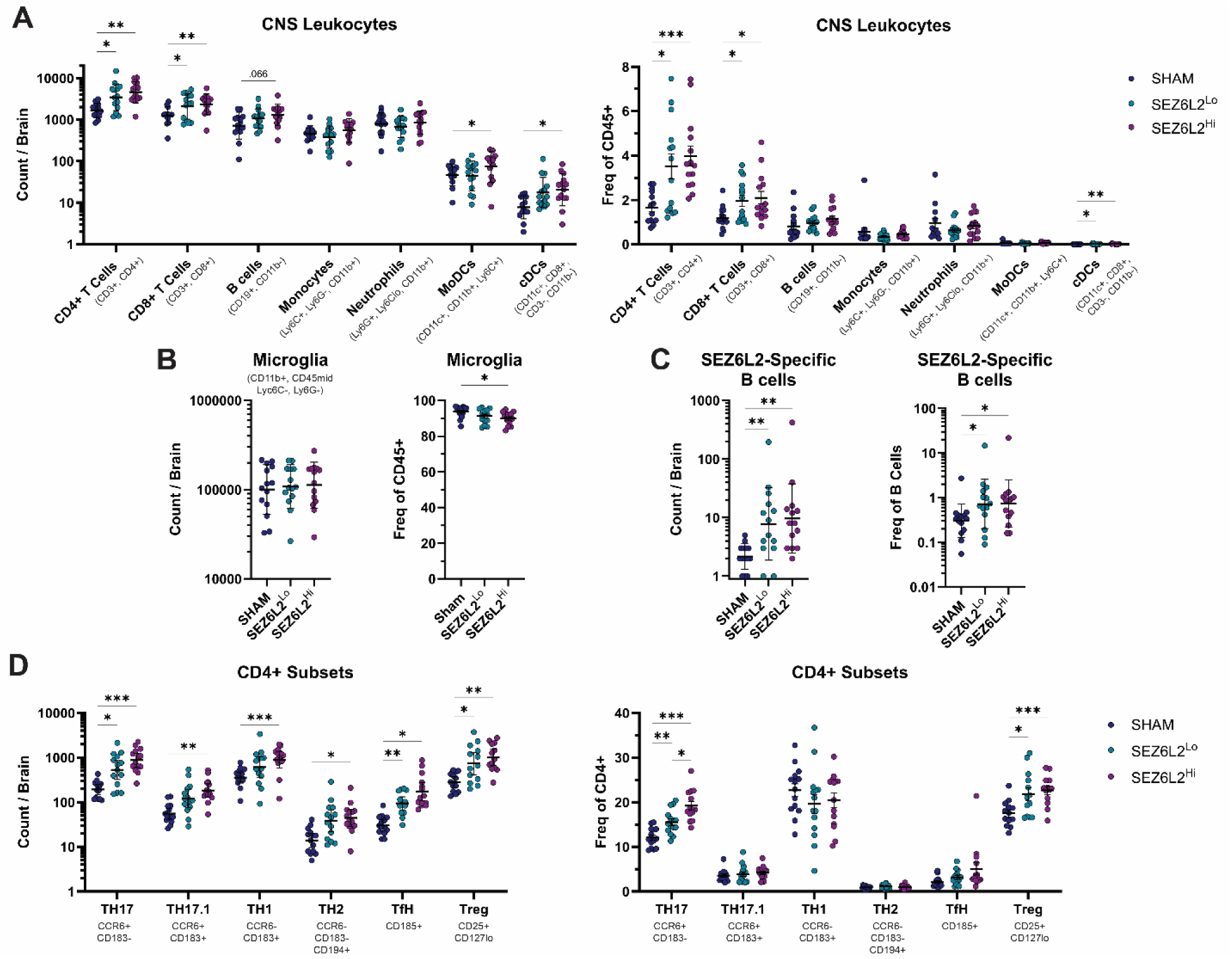
SJL mice show expanded CNS immune infiltration following SEZ6L2 immunization. Flow cytometric analysis of CD45⁺ immune cell populations in the brain at 6 weeks post-immunization. **A-B)** Absolute counts (left) and frequencies within CD45⁺ cells (right) for the indicated immune populations. Statistics: repeated-measures 2-Way ANOVA with Tukey’s multiple comparisons test. **C)** SEZ6L2-specific B cells identified by binding to fluorescently labeled recombinant SEZ6L2, shown as absolute counts (left) and frequency within CD19⁺ B cells (right). Statistics: Kruskal-Wallis Test with Dunn’s multiple comparisons test. **D)** CD4⁺ T-cell subsets in the CNS, shown as absolute counts (left) and frequency within CD4⁺ T cells (right). Statistics: Mixed-effects model with Tukey’s multiple comparisons test. For all graphs: Data are presented as geometric means +/- SD; n=14 mice per group.

### Identification of SEZ6L2-specific B cells within the CNS

Given the robust antibody responses observed in SJL mice, we next asked whether antigen-specific B cells could be directly detected within the CNS. Using fluorescently labeled recombinant human SEZ6L2 protein, we identified a discrete population of SEZ6L2-binding CD19⁺ B cells in the brains of immunized mice that were largely absent in sham controls (Figure 4C). These SEZ6L2-specific B cells could potentially enable local antibody production in the CNS.

### Increase of multiple CD4⁺ T-cell subsets in the CNS

To further define the composition of the CD4⁺ T-cell response, we analyzed subtype distributions using chemokine receptor and surface marker expression. CD4⁺ T-cell subsets were defined as follows: TH17 (CD183⁻, CCR6⁺), TH17.1 (CD183⁺, CCR6⁺), TH1 (CD183⁺, CCR6⁻), TH2 (CD183⁻, CCR6⁻, CD194⁺), T follicular helper (TfH; CD185⁺), and regulatory T cells (Treg; CD25⁺, CD127^lo^). SEZ6L2 immunization resulted in significant increases in the absolute number of all CD4⁺ T-cell subsets within the CNS (Figure 4D). However, when analyzed as a proportion of total CD4⁺ T cells, only TH17 and Treg populations showed significant increases in frequency. These findings show that while SEZ6L2 immunization drives broad expansion of CD4⁺ T-cell subsets in the CNS, selective enrichment of TH17 and regulatory populations further compete to tune the overall immune response.

### SEZ6L2 immunization induces behavioral deficits associated with coordinated immune responses

To determine whether amplified immune responses in SJL mice were associated with neurological dysfunction, we performed a battery of behavioral assays. Baseline performance in the open field and rotarod assays confirmed comparable pre-immunization motor function across groups (Supplemental Figure 2). At 2.5 weeks post-immunization, SEZ6L2-immunized SJL mice exhibited increased locomotor activity in the open field compared to sham controls (Figure 5A). This early hyperactivity resolved by 5.5 weeks, at which point total distance traveled was no longer different between groups. At 5.5 weeks, immunized mice also exhibited increased foot faults on both the wire grid and tapered beam assays, as well as increased time to descend in the pole test (Figure 5B–E). A trend toward reduced latency to fall on the accelerating rotarod was also observed (SHAM vs SEZ6L2^Hi^; p=0.074). These motor deficits were generally more pronounced in the high-dose group, although differences between dose groups were mostly insignificant.

**Figure 5:**
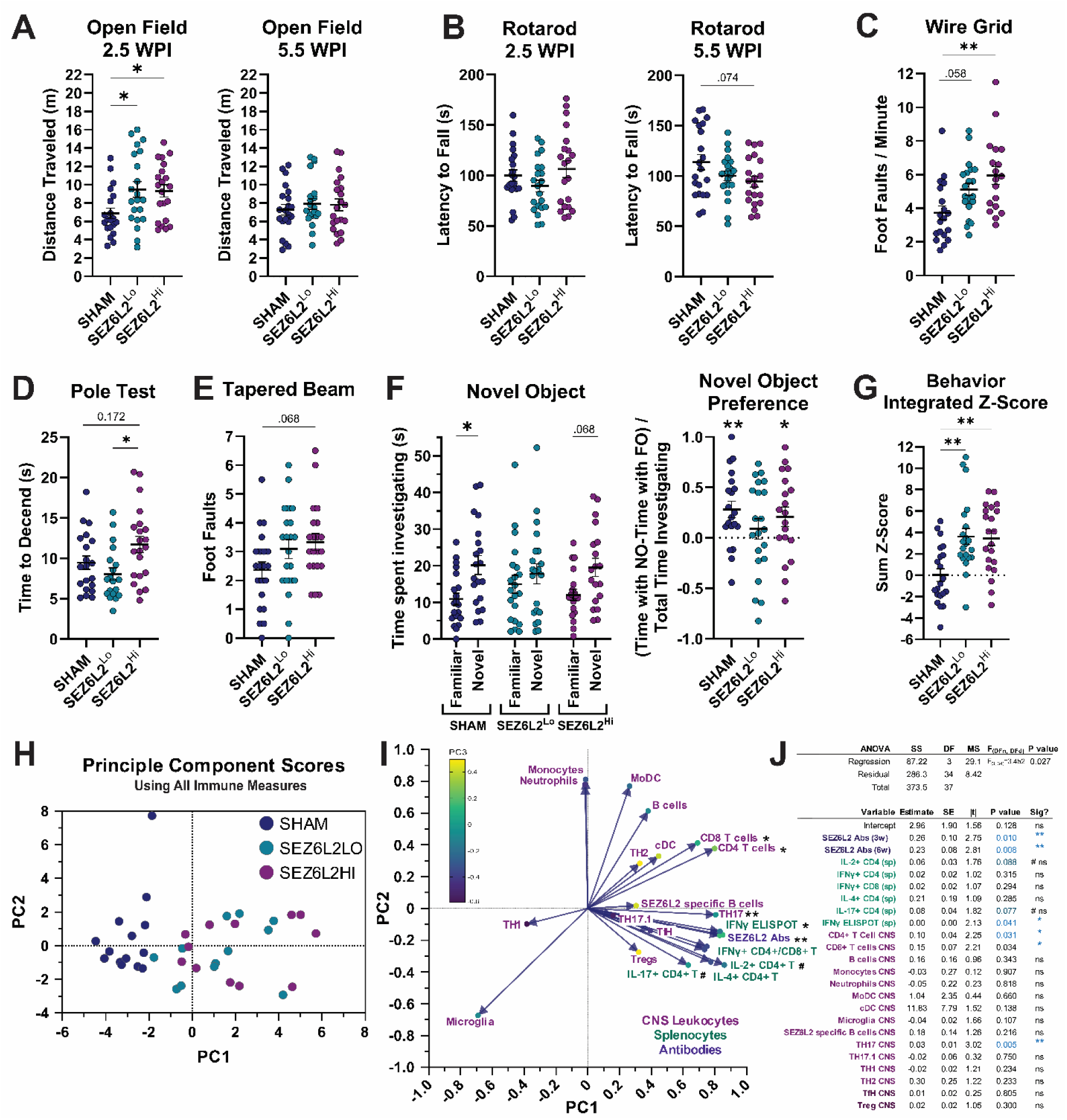
SEZ6L2 immunization induces behavioral deficits associated with coordinated immune responses in SJL mice. **A)** Distance traveled in the open field assay at 2.5- and 5.5-weeks post immunization (WPI). **B)** Latency to fall from an accelerating rotarod at 2.5 and 5.5 WPI. **C)** Foot faults on a wire grid normalized to minutes of active movement at 5.5 WPI. **D)** Time to descend in the pole test at 5.5 WPI. **E)** Foot faults on the tapered beam assay at 5.5 WPI. **F)** Novel object recognition assay at 5.5 WPI. Left: time spent investigating familiar vs. novel objects. Right: Novel object preference index (time spent with the novel object normalized to total object investigation time). **G)** Integrated behavioral Z-score calculated as the sum of standardized z-scores from the 2.5 WPI open field distance and all 5.5 WPI behavioral assays. **H-J)** Principal component analysis (PCA) of 23 immune variables derived from figures 2-4. **H)** Principal component score plot showing separation of SEZ6L2-immunized mice from SHAM controls (each point represents one mouse). **I)** Loadings plot illustrates the contribution of individual immune variables to each principal component. **J)** Principal component regression using the three PCs to predict the integrated behavioral Z-score. **Statistical analysis** for A-G: 1-way ANOVA with Tukey’s multiple comparison test; Mean +/- SEM; n=18-21 mice per group. For the Novel Object Preference graph in F, a one sample t-test was used relative to a theoretical mean of 0. Principal component regression (least squares) significantly predicted behavioral Z-scores (p = 0.0271; F(3,34) = 3.452; R² = 0.2335). Asterisks (*) indicate p < 0.05 and (**) p < 0.01; (#) indicates p < 0.1.

To assess potential cognitive effects, we performed a novel object recognition assay. Sham mice displayed the expected significant preference for the novel object over the familiar object, while the high-dose SEZ6L2-immunized mice showed a similar, but insignificant trend. In contrast, mice in the low-dose SEZ6L2 immunization group failed to show this preference, suggesting a deficit in recognition memory (Figure 5F).

Given the variability inherent in individual behavioral assays, we next calculated an integrated behavioral score to capture overall functional impairment. For each mouse, Z-scores from the 2.5 WPI open field distance and all 5.5 WPI behavioral assays were summed to generate a composite behavioral metric. Both low- and high-dose SEZ6L2-immunized groups showed significant impairment relative to sham controls, with no significant difference between dose groups (Figure 5G). Together, these findings show that SEZ6L2 immunization induces moderate motor deficits consistent with cerebellar dysfunction, accompanied by variable effects on cognitive performance that may reflect contributions from cerebellar and extra-cerebellar SEZ6L2-expressing circuits.

To determine whether immune responses were associated with behavioral outcomes, we performed principal component analysis (PCA) using 23 immune variables, including serum antibody levels, splenocyte responses to SEZ6L2 stimulation, and CNS leukocyte populations. PCA identified three principal components that captured variance beyond that expected by parallel analysis of “random” data (PC1 = 35.7%, PC2 = 16.8%, PC3 = 10.9% variance). Visualization of mice in principal component space revealed clear separation between SEZ6L2-immunized and sham groups (Figure 5H), while low- and high-dose groups remained largely overlapping, consistent with similarities observed in individual immune measures.

We next used these principal components as predictors in a regression model to assess their relationship with behavioral impairment, as defined by the integrated behavioral Z-score. Principal component regression revealed a significant association between immune features and behavioral outcomes (p = 0.0271; F(3,34) = 3.452), explaining approximately 23% of the variance in behavioral scores (R² = 0.2335; Figure 5J). Immune variables contributing most significantly to the model included SEZ6L2 antibody levels at 3 and 6 weeks, IFNγ ELISPOT responses following SEZ6L2 stimulation, and CNS frequencies of CD4⁺ T cells, CD8⁺ T cells, and TH17 cells. These findings support a quantitative relationship between coordinated humoral and cellular immune responses and functional neurological impairment in this model.

## DISCUSSION

This study demonstrates that genetic background influences both the magnitude and organization of SEZ6L2-directed immune responses in mice. While immunization induces coordinated humoral and CNS immune activation across strains, SJL mice develop an earlier and more robust response, including heightened antibody production and expanded infiltration of T and B lymphocytes into the CNS. This amplified immune response is associated with measurable neurological dysfunction. SEZ6L2-immunized SJL mice develop moderate but reproducible motor deficits consistent with cerebellar impairment, along with more variable effects on cognitive performance. When behavioral outcomes were integrated across assays, both immunized groups showed a consistent overall deficit. In addition, multivariate analysis demonstrated that combined immune features were significantly associated with behavioral outcomes, supporting a link between coordinated immune responses and neurological dysfunction. These findings establish SJL mice as an enhanced model of SEZ6L2 autoimmunity.

Compared to our previously published study with C57BL/6 mice^17^, this study looked more extensively at the immune cell populations infiltrating the CNS. In particular, we were able to detect SEZ6L2-specific B cells in the brain, which implies that they may have local roles in anti-SEZ6L2 antibody production, antigen presentation, and interactions with T-cells. While the C57BL/6 model only showed an increase in CD4+ T cells, the SJL model showed that both CD4+ and CD8+ T cell populations were significantly increased. Thus, the SJL model is likely more prone to neuropathology caused by SEZ6L2-specific cytotoxic T cells than the C57BL/6 model. Additionally, the increased representation of TH17 and TH17.1 CD4⁺ T-cell subsets in the CNS may further potentiate pathology, as these populations are known to promote blood–brain barrier disruption, recruit additional immune cells, and amplify neuroinflammation in models such as experimental autoimmune encephalomyelitis.^40, 41^ TH17 cells also support local B-cell activation (including class switching, antibody production, and germinal center-like responses in meningeal ectopic lymphoid tissue), amplifying humoral immunity and B-cell antigen presentation within the CNS.^42, 43^ The enhanced Treg populations also show that self-tolerance mechanisms are actively working to suppress the immune response to SEZ6L2 as an effort to protect the CNS. How the balance of Treg suppression and TH17 activation plays out in each individual mouse likely contributes to the variability in neurological phenotypes.

The immunodominant T-cell epitopes identified in SJL mice differed from those observed in C57BL/6 mice^17^, consistent with differences in MHC class II haplotypes and reinforcing the role of host genetics in shaping antigen presentation and T-cell specificity. In humans, SEZ6L2-directed autoimmune responses are also expected to be influenced by genetic context and to vary across HLA haplotypes. Consistent with this, patients with idiopathic/sporadic ataxias show increased rates of autoimmune disease and enrichment of specific HLA alleles, including HLA-DQ2^44^, supporting the importance of evaluating HLA associations in future cases of SEZ6L2 autoimmunity. The mouse epitopes identified here also provide experimental tools for future studies, enabling the development of peptide–MHC class II tetramers, peptide-based T-cell assays, and peptide immunization strategies to more precisely interrogate antigen-specific responses.

Interestingly, behavioral outcomes did not clearly scale with immunization dose. This suggests that once a threshold level of immune activation is reached, additional increases may not translate into greater functional impairment. It is also possible that factors such as the localization of immune activity or target engagement within the brain contribute to variability in behavioral outcomes.

Several important questions remain. It is still unclear whether SEZ6L2 autoantibodies are directly pathogenic, which epitopes drive neuronal dysfunction, and how antibody and cellular immune responses interact within the CNS. The SJL model provides a platform to address these questions through direct functional analysis of autoantibodies and antigen-specific lymphocytes.

Together, the SJL and C57BL/6 models provide complementary systems for studying SEZ6L2 autoimmunity. The reproducibility of key features across strains supports the biological relevance of this model, while the enhanced immune response in SJL mice may enable more detailed mechanistic interrogation. These models provide a foundation for defining pathogenic pathways and developing targeted therapeutic strategies for SEZ6L2-associated neurological disease.

## Availability of data and materials

The datasets used and/or analyzed during the current study are available from the corresponding author on reasonable request.

## Authors’ contributions

J.H. designed the study. C.R.S., J.R., and J.H. performed the immunizations. All authors contributed to animal care, as well as end-point tissue collection and processing. C.R.S. and J.H. conducted the behavioral experiments. C.R.S. performed the SEZ6L2-specific T-cell flow cytometry assays and antibody ELISAs. J.G., with assistance from A.H., C.R.S., and J.H., conducted the ELISPOT assays. J.R. performed flow cytometry analysis of brain samples. All authors reviewed and approved the final manuscript.

## Acknowledgements

We thank Dr. Kerry O’Banion and Dr. Christoph Proschel for generously providing access to their behavior equipment.

## Supplementary Material

**Supplemental Figure 1:**
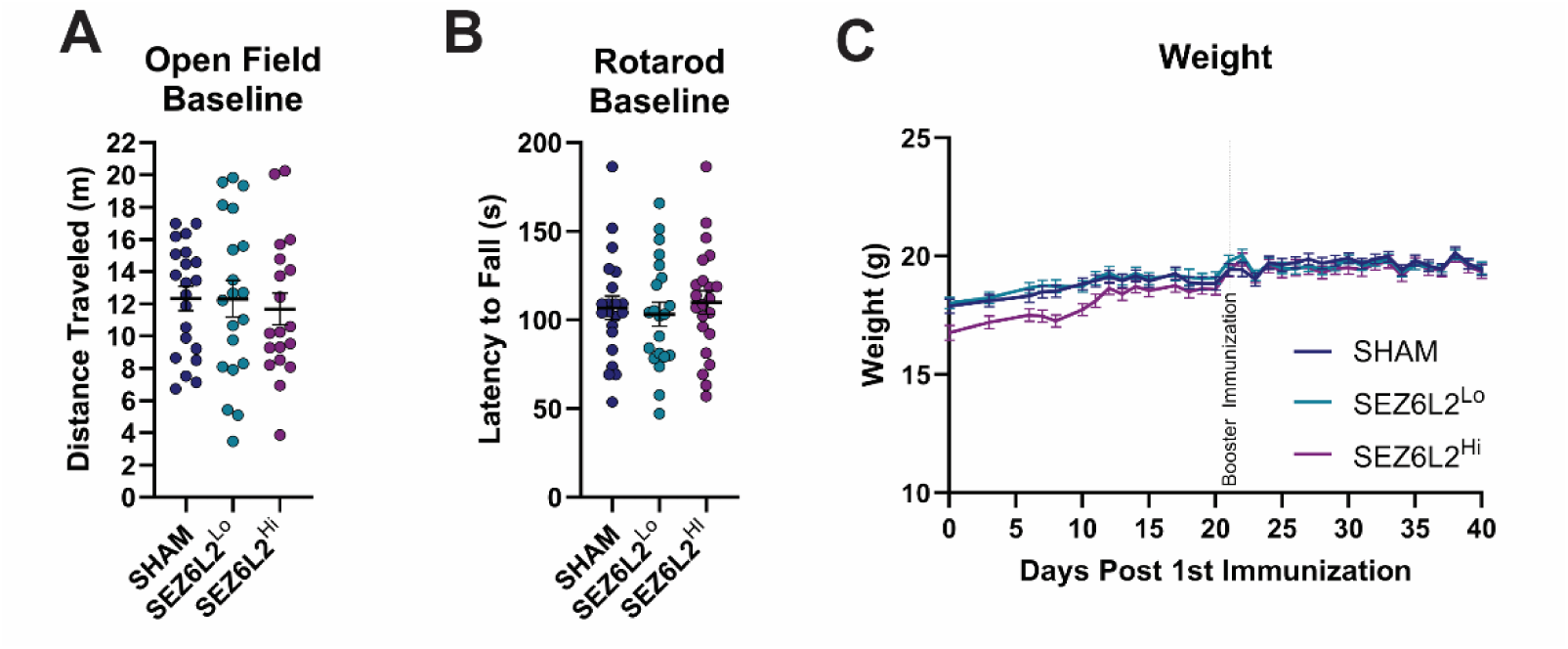
SJL experimental groups showed no baseline behavioral differences prior to immunization and no significant changes in weight following immunization. **A)** Distance traveled in an open field assay prior to immunization. **B)** Latency to fall on the rotarod prior to immunization. **C)** Body weights measured over 6 weeks post-immunization. Random group assignment resulted in the SEZ6L2 high-dose group having a lower average baseline weight compared to other groups; however, weights equalized over time, and no group exhibited significant weight loss following priming or booster immunizations. Statistical analysis for A and B: 1-way ANOVA with Tukey’s multiple comparison test; Mean +/- SEM; n=21 mice per group.

**Supplemental Figure 2:**
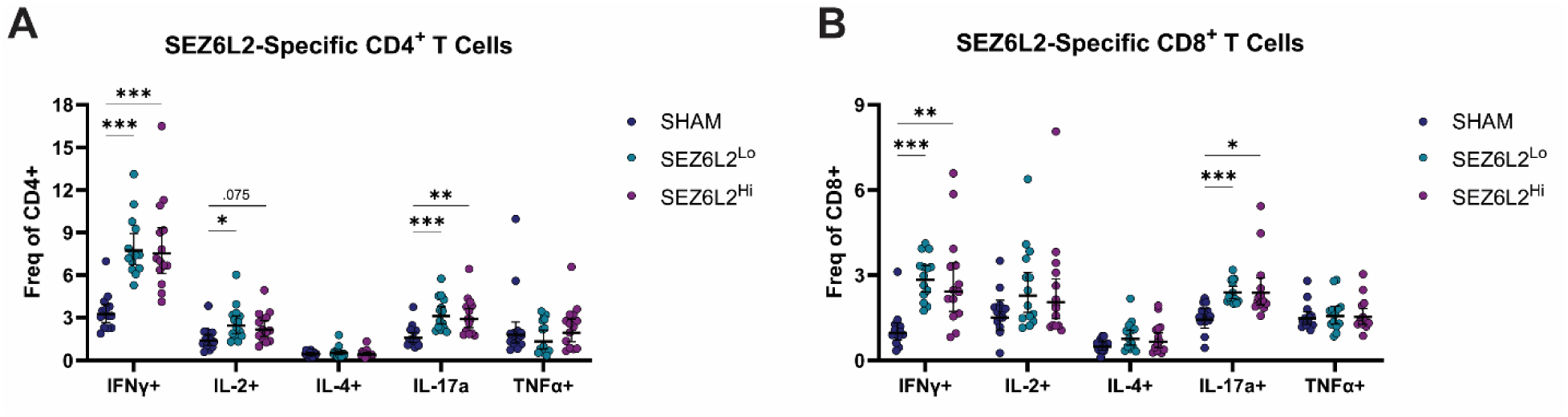
SEZ6L2-Specific CD4+ and CD8+ T Cells. Flow cytometric analysis of CD4⁺ **(A)** and CD8⁺ **(B)** T cell populations for intracellular expression of the indicated cytokines after 72-hour stimulation with recombinant SEZ6L2. Statistics: repeated-measures 2-Way ANOVA with Geisser-Greenhouse correction and Tukey’s multiple comparisons test; Data are presented as means +/- SEM; n=14 mice per group.

**Supplemental Figure 3:**
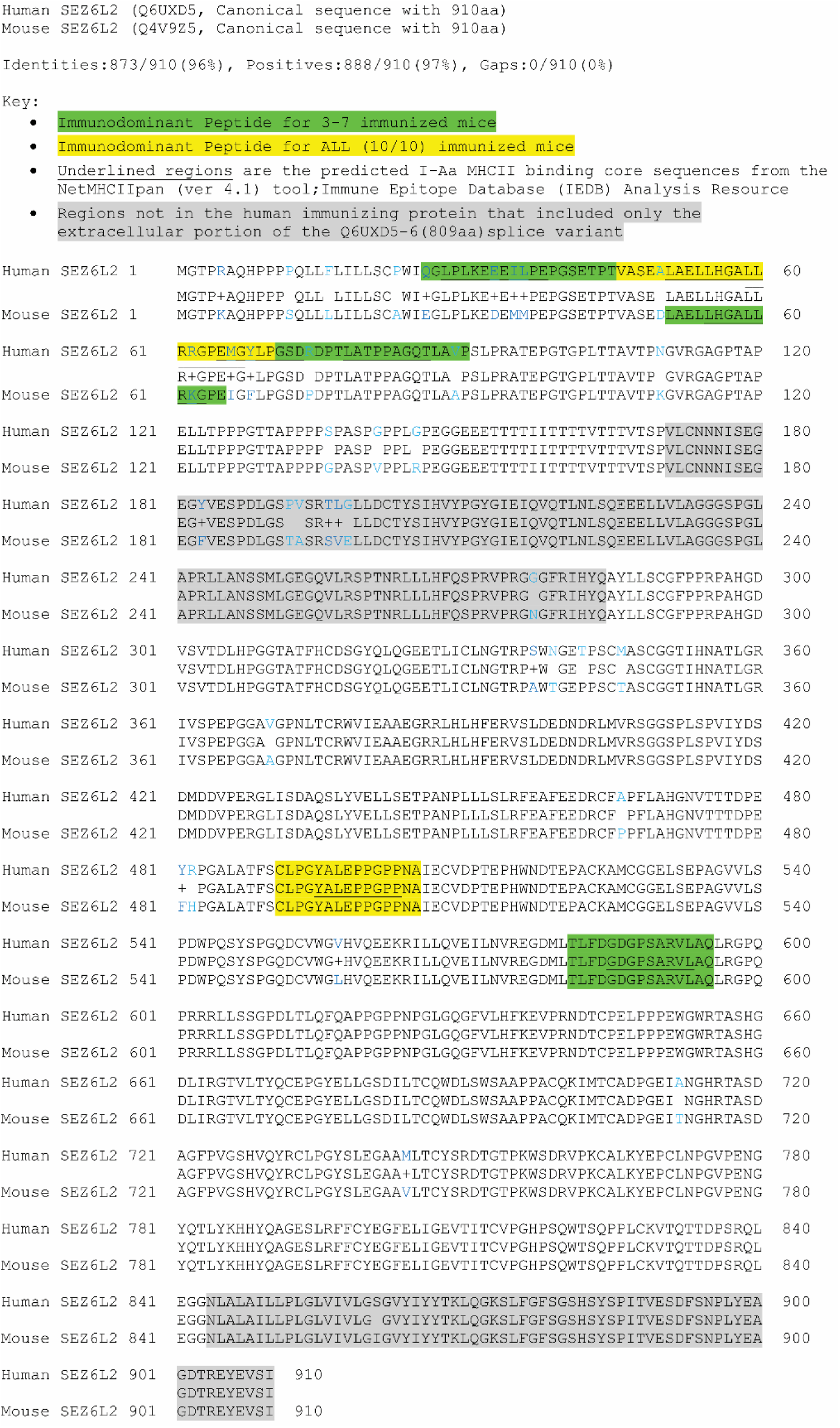
Protein BLAST alignment of human and mouse SEZ6L2 sequences with immunodominant peptides underlined and highlighted as indicated.

